# Visiomode: an open-source platform for building rodent touchscreen-based behavioral assays

**DOI:** 10.1101/2022.12.01.518732

**Authors:** Constantinos Eleftheriou, Thomas Clarke, Victoriana Poon, Marie Zechner, Ian Duguid

**Affiliations:** Simons Initiative for the Developing Brain, University of Edinburgh, Edinburgh, EH8 9XD, UK; Centre for Discovery Brain Sciences and Patrick Wild Centre, Edinburgh Medical School: Biomedical Sciences, University of Edinburgh, Edinburgh, EH8 9XD, UK; The Roslin Institute, University of Edinburgh, Easter Bush, Midlothian, EH25 9RG, Scotland, UK

## Abstract

**Background:** Touchscreen-based behavioral assays provide a robust method for assessing cognitive behavior in rodents, offering great flexibility and translational potential. The development of touchscreen assays presents a significant programming and mechanical engineering challenge, where commercial solutions can be prohibitively expensive and open-source solutions are underdeveloped, with limited adaptability.

**New method:** Here, we present Visiomode (www.visiomode.org), an open-source platform for building rodent touchscreen-based behavioral tasks. Visiomode leverages the inherent flexibility of touchscreens to offer a simple yet adaptable software and hardware platform. The platform is built on the Raspberry Pi computer combining a web-based interface and powerful plug-in system with an operant chamber that can be adapted to generate a wide range of behavioral tasks.

**Results:** As a proof of concept, we use Visiomode to build both simple stimulus-response and more complex visual discrimination tasks, showing that mice display rapid sensorimotor learning including switching between different motor responses (i.e., nose poke versus reaching).

**Comparison with existing methods:** Commercial solutions are the ‘go to’ for rodent touchscreen behaviors, but the associated costs can be prohibitive, limiting their uptake by the wider neuroscience community. While several open-source solutions have been developed, efforts so far have focused on reducing the cost, rather than promoting ease of use and adaptability. Visiomode addresses these unmet needs providing a low-cost, extensible platform for creating touchscreen tasks.

**Conclusions:** Developing an open-source, rapidly scalable and low-cost platform for building touchscreen-based behavioral assays should increase uptake across the science community and accelerate the investigation of cognition, decision-making and sensorimotor behaviors both in health and disease.

## Introduction

Since their introduction to biomedical research, touchscreens have become an increasingly popular tool for assessing cognitive function in rodents (Bussey et al., 1997; Dumont et al., 2021; Markham et al., 1996). Their appeal lies with their remarkable flexibility, supporting a vast array of visual stimuli coupled with quantifiable motor responses (Seitz et al., 2021), both of which are necessary for designing tasks to investigate complex cognitive processes such as category learning (Broschard et al., 2021; Kim et al., 2018), spatial attention (Haddad et al., 2021), cognitive flexibility (Groman et al., 2012), and visual perception (Markham et al., 1996). Their use has transformed studies of neurological disorders by providing a sensitive assay of sensorimotor behaviors (Arulsamy et al., 2019; Copping et al., 2017; Leach and Crawley, 2018; Leach et al., 2016; Morton et al., 2006; Norris et al., 2019; Yang et al., 2015), revealing subtle phenotypes that were not detected by more conventional assays (Van den Broeck et al., 2019; Zeleznikow-Johnston et al., 2018). This sensitive readout of changes in behavior holds great translational promise (Talpos and Steckler, 2013), where tasks designed for animals can be directly translated to human subjects (Chow et al., 2020; Hvoslef-Eide et al., 2015; Nithianantharajah et al., 2015). Despite their increasing popularity, the use of touchscreen-based behaviors is somewhat limited in rodent research. Uptake has been hampered either by the prohibitive up-front costs of commercial systems or the considerable ‘in-house’ development required to create bespoke touchscreen-based behaviors (Dumont et al., 2021).

Commercially available touchscreen behavioral arenas provide researchers with a simple turnkey solution requiring minimal setup time. These systems have dominated the touchscreen landscape in biomedicine over the past two decades (Arulsamy et al., 2019; Brasted et al., 2002; Brigman et al., 2010; Bussey et al., 1998; Bussey et al., 2008; Delotterie et al., 2014; Glover et al., 2020; Haddad et al., 2021; Heath et al., 2019; Odland et al., 2021; Piantadosi et al., 2019; Stirman et al., 2016; Talpos et al., 2008), and have played an important role in popularizing their use in rodent research (Dumont et al., 2021). However, the prohibitive costs associated with commercial systems (i.e., > 10,000 USD) provides a rate limiting step for their widespread adoption (Dumont et al., 2021). In contrast, developing touchscreen tasks ‘inhouse’ is a particularly challenging programming and engineering problem. While most traditional open-field (Hall and Ballachey, 1932) or operant chamber (Skinner, 1938) tasks can be implemented with a simple microcontroller device (Akam et al., 2022), the introduction of a touchscreen interface requires complex hardware and software integration to control the generation and display of graphics, as well as registering behavioral interactions with the screen. Utilizing graphics libraries available on most Operating Systems (e.g., Microsoft Windows, Linux and MacOS) requires extensive programming knowledge (Kessenich et al., 2016), and while open-source initiatives such as PsychoPy greatly simplify the task of generating visual stimuli (Peirce, 2007), they still require significant development to be adapted for touchscreen tasks (Seitz et al., 2021). This is further complicated by the choice of touchscreen hardware, where heterogeneity in compact touchscreen systems results in variable touch sensitivities, requiring the developer to test and validate a range of screens before final implementation (Dumont et al., 2021).

In our view, an open-source, community-driven touchscreen solution would solve both problems by distributing the development effort across multiple research groups, while also reducing overall costs (Fortunato and Galassi, 2021; Freeman, 2015). Open-science initiatives have continued to grow in the past few years, with projects like Mouse-Bytes (Beraldo et al., 2019) and the advice sharing platform touchscreencognition.org (Dumont et al., 2021). To date, no open-source community-based solution exists. While open-source touchscreen-based operant chambers have been developed (Gurley, 2019; O’Leary et al., 2018; Pineno, 2014), this has not led to increased uptake due a lack of code availability and the primary focus being on reducing costs, rather than enhancing the user experience, scalability, adaptability, and ease of use.

To address these unmet needs, we have developed Visiomode (www.visiomode.org), a complete open-source software and hardware platform for developing touchscreen-based behavioral tasks for rodents. Visiomode combines a sophisticated web-based user interface (UI) and powerful plug-in system, with an affordable hardware configuration that can accommodate a wide variety of visuomotor tasks (Figure 1). Our solution has been designed to maximize adaptability, while providing a consistent, standardized user experience and output data format. While several visual stimuli and task structures have been provided out-of-the-box, including drifting gratings, symbols and natural images, users can upload or simply programmatically define their own visual stimuli. Using simple stimulus-response and more complex visual discrimination tasks as exemplars, we show that mice display rapid sensorimotor learning, switching between both nose poke and visually guided reaching depending on task requirements. In addition, we discuss Visiomode’s Application Programming Interface (API), how it can be scaled to parallelize data acquisition using a single personal device and the build components.

**Figure 1.**
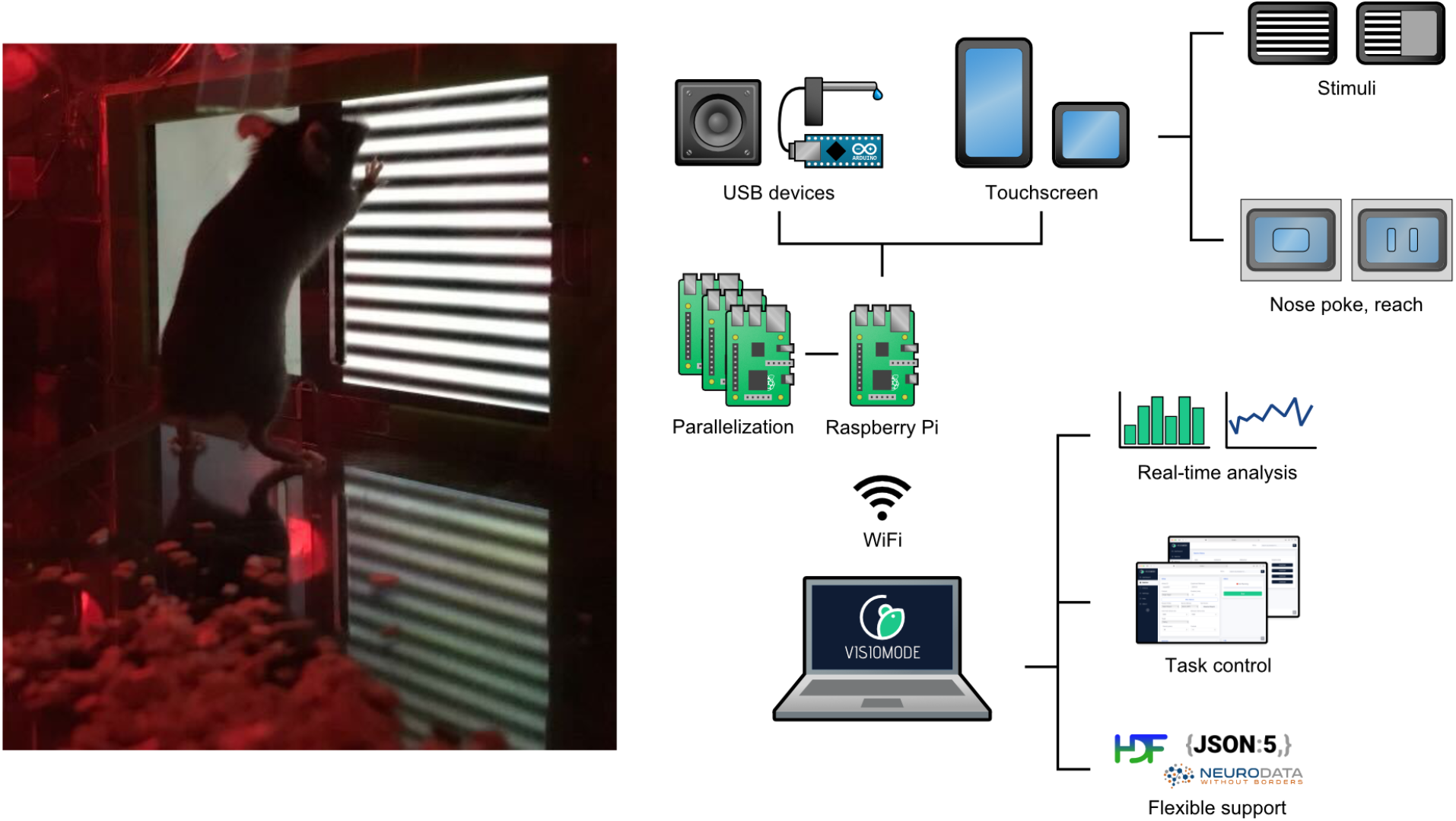
Visiomode: a flexible, scalable platform for building touchscreen-based tasks. *Left*, Image of the mouse behavioral arena with interactive touchscreen controlled by Visiomode. *Right*, Visiomode is a web-based interface that controls, via WiFi, a Raspberry Pi computer and any coupled USB devices (e.g. loudspeaker, USB microcontroller for reward delivery, touchscreen). Visual stimuli can be generated programmatically via Visiomode’s API or loaded as images / animation files on-the-fly and presented via any touchscreen supported by Raspberry Pi, including integrated touchscreen displays such as the Pimoroni Hyperpixel. Visiomode also supports real-time analysis with data being exported in a variety of formats, including JSON, HDF5 and NWB.

## Methods

### API design principles

We first designed Visiomode’s API which encapsulates all the software functionality required to design and run touchscreen behavioral tasks, including stimulus generation, trial structure definition, response recording and interfacing with external hardware via the USB. The API is written in Python, leveraging its popularity, wide availability of libraries and ease of use. The PyGame and PySDL libraries are used for handling graphics and task timing, while the Flask library is used for rendering Visiomode’s web interface. In addition to the behavioral tasks provided out-of-the-box, the API can be used to implement additional task components such as protocols, stimuli or hardware integrations using user-defined task files.

The API is broadly divided into three parts: stimulus interface, protocol interface and device interface. Each constituent part represents a Python abstract base class (Hunt, 2019), which defines a programmatic interface which all user-defined stimuli, protocols and devices must be derived from. Each interface allows users to integrate custom HTML forms for dynamically setting component-specific parameters within Visiomode’s web interface.

The stimulus interface allows for user-defined visual stimuli to be integrated into behavioral tasks, independently from task structure. For example, the same user-defined stimulus, such as a solid color or a drifting grating, can be reused across all available task protocols without having to be redefined. Each class derived from the stimulus interface inherits several functions that are core to the presentation of the visual stimuli within a protocol, such as functions that control the appearance and removal of stimuli, updating of stimulus position, or modification of stimuli between trials. Additionally, stimuli can integrate peripheral USB devices to yield multi-sensory stimuli (i.e., simultaneous presentation of an auditory tone and drifting grating).

Task structure definition classes are inherited from the protocol interface, which controls the timing for the presentation of stimuli and processes touchscreen and external device events during an experimental session. User-defined tasks must specify, as a minimum, a target stimulus parameter, which may be set dynamically via the web interface. The web interface will pass the session duration, inter-trial interval (ITI) and duration of stimulus presentation to every protocol-derived class, which by default will iterate through calls to a trial_block function until the session duration expires. The trial_block function monitors the application’s touch event queue during the ITI and while a stimulus is present, and assigns correct, incorrect, uncued or miss responses accordingly. The protocol interface implements several functions corresponding to different trial outcome conditions, such as on_ correct, on_incorrect and on_uncued, as well as trial epochs such as on_trial_start or on_ stimulus_start, which can be overridden by users to build complex trial structures with fewer lines of code and without requiring an extensive understanding of Visiomode’s core functionality.

While most behavioral protocols will be defined before each session, Visiomode can support dynamic, on-the-fly changes to stimulus and protocol settings depending on performance. For example, the contrast of a stimulus could be decreased following a user-defined number of correct responses within a single training session. This would be achieved by implementing a new protocol file, which can then be uploaded to Visiomode via the web interface. For more information, we encourage users to visit www.visiomode.org for the latest guidance on implementing new protocols.

While Visiomode’s web interface can be accessed via a browser, instances of Visiomode do not need an active internet connection as they contain all necessary code libraries for running tasks and monitoring performance, including stimulus generation and live plotting of behavioral data. While the recommended installation route requires an internet connection, we provide alternative means of installing the software on Raspberry Pis if internet connectivity is not possible.

Finally, in addition to the touchscreen itself, Visiomode can integrate external USB devices connected to the Raspberry Pi through the device interface. For example, a water reward mechanism driven by an Arduino microcontroller can be used to dispense rewards following correct task responses. The device interface is subdivided into Input and Output interfaces, supporting both sensors that can feed into a task structure as well as actuators providing additional sensory stimuli or dispensing rewards. While the Raspberry Pi’s own GPIO ports could also be used to integrate sensors or actuators in Visiomode tasks, supporting USB devices can be advantageous. Touchscreen displays for the Raspberry Pi use all the available GPIO ports, which leads to a more compact design, but necessitates the use of USB-connected microcontrollers to integrate external hardware. Given the ubiquity of microcontroller devices in neuroscience, this plug-and-play approach allows the user to easily incorporate USB-connected devices from existing setups facilitating friction-free development of novel behavioral tasks.

While Visiomode has been designed as a flexible platform that can be readily extended programmatically, it offers several common experimental paradigms out-of-the-box providing utility for users with no prior programming experience. Single target, two-alternative forced choice (2AFC) and Go/NoGo protocols are included with every installation which support correction trials, pseudo-randomization of stimulus presentation, and any arbitrary external reward devices. Sinusoidal gratings with user-defined characteristics, as well as a range of symbols and natural scenes are also included, which can be incorporated into any protocol. Finally, Visiomode supports a range of external reward devices, such as reward spouts and food hoppers, with microcontroller code compatible with Visiomode available at https://doi.org/10.5281/zen-odo.6877795. For any USB connected devices for which the Rasberry Pi operating systems does not provide an out-of-the-box solution for integration/calibration, Visiomode supports the use of custom written “driver” code within its devices interface.

To facilitate synchronization of behavior and physiological recordings, Visiomode time-stamps behavioral epochs using the system clock, which can be synched with a local Network Time Protocol server to provide sub-millisecond time synchronization between the Raspberry Pi computer running Visiomode and the computer acquiring physiological data. For physiological recordings that require higher temporal precision (i.e., electrophysiological recordings at 20 KHz), Visiomode can provide a time synchronization signal using the host Raspberry Pi’s GPIO ports or a microcontroller device connected to the USB (Akam et al., 2022).

Visiomode’s source code is openly available and is publicly hosted at https://doi.org/10.5281/zenodo.6877795 under the terms of the MIT license.

### Web interface

Each instance of Visiomode exposes a web interface that enables the user to set up, run and monitor touchscreen experiments (Figure 2), accessible via any web browser that is connected to the same local network as the Raspberry Pi. The web interface is designed to cater to both novice and expert users. An accessible user interface enables the uptake of Visiomode by users with no prior programming knowledge, while offering a fully customizable user interface for users that wish to extend Visiomode’s capabilities. The primary function of the web interface is to set up and run touchscreen behavioral tasks. Task, stimulus, and device parameters can be set on-the-fly, allowing users to control all aspects of the setup without the need to modify the code. For example, Visiomode provides out-of-the-box support for drifting gratings, where parameters such as the cycles per degree, contrast and drift frequency can be adjusted online. Each component, derived from the Protocol, Stimulus and Device interfaces described in the previous section, can optionally implement a webform via the form_path attribute that passes parameters from the web interface to the Visiomode API (for an example, see https://doi.org/10.5281/zenodo.6877795) (Figure 2). These webforms are loaded dynamically when the user selects a particular component and can interface with multiple components to build complex tasks (e.g., protocols can have multiple stimuli integrated with multiple input and/or output devices). The parameters set on the web interface are serialized and converted to asynchronous calls to the Visiomode API, which assembles the different components into a behavioral protocol that runs on the touchscreen.

**Figure 2.**
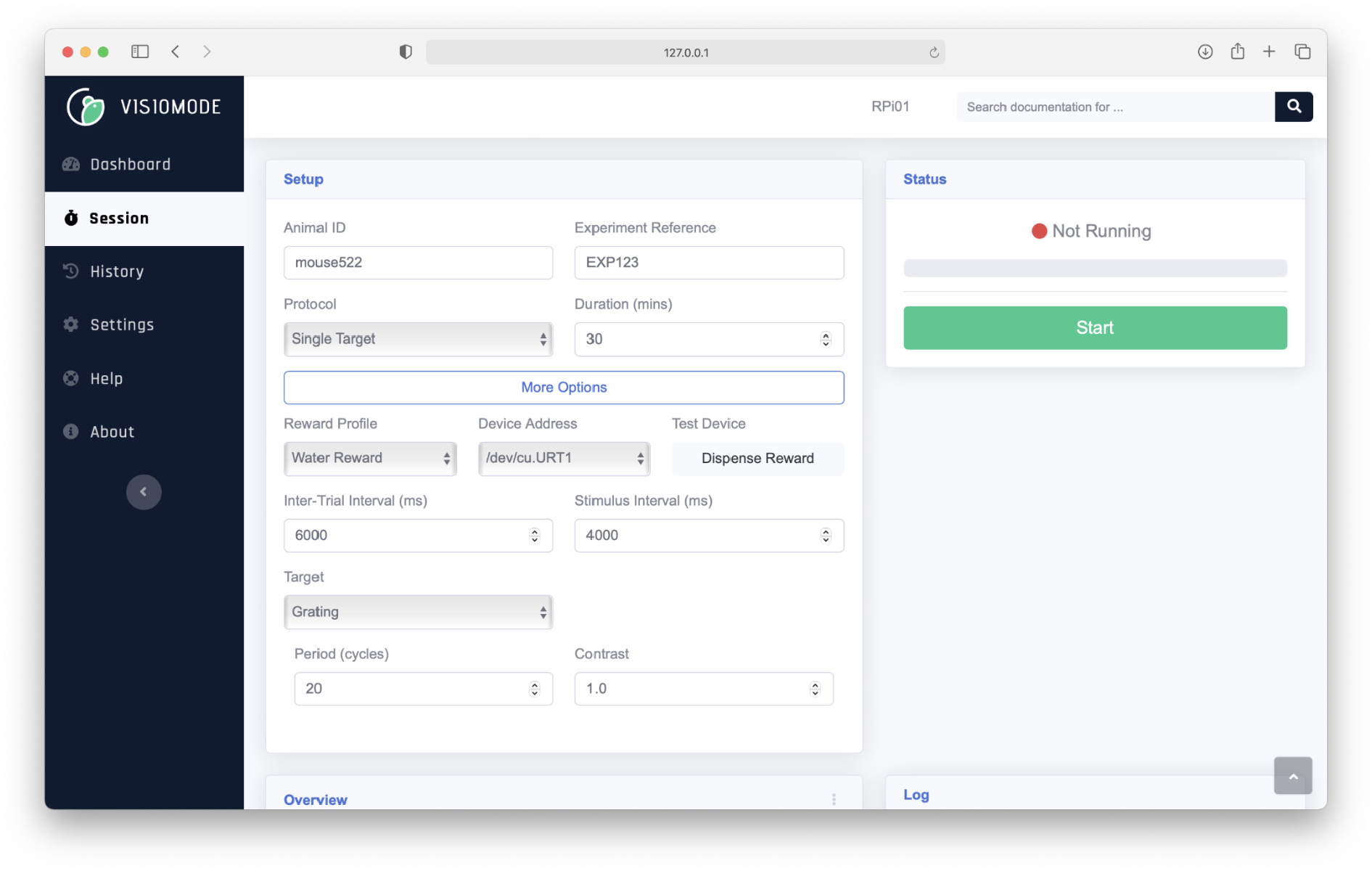
Visiomode: web-based Graphical User Interface (GUI). Visiomode GUI which provides session information for an individual mouse (mouse552) and experiment (single drifting grating target, nose poke). The ‘More Options’ tab allows the user to define advanced stimulus parameter options such as frequency, contrast, inter-trial-interval duration and separator size (pixels).

In addition to task control, the Visiomode interface plots real-time task analytics that are protocol-specific and can be further customized or extended via an analytics_path attribute, see https://doi.org/10.5281/zenodo.6877795. Visiomode analytics use the Graphs.js Javascript library for plotting, however other popular Javascript graphics libraries such as D3 or Plotly can also be used. Visiomode’s web interface pools data from the API running the behavior at 4 second intervals, updating only the analytics components visible to the user.

### A flexible, scalable platform for building touchscreen tasks

The web interface design ensures that each Visiomode setup is self-contained, and that there is virtually no upper limit to parallelization, providing all devices are connected to the same local network. Each instance of the web interface is designed to be asynchronous to the running of the task, such that if the web interface is disconnected (e.g., by an intermittent network issue, or accidental closing of the device’s browser) the Visiomode API running the task will be unaffected. Decoupling the web interface from the API running the experimental session yields a resilient and easily scalable core platform, which users can customize and extend to suit their individual experimental needs.

The web interface can also be used to export session data in a variety of different formats. By default, Visiomode session data are stored as human-readable Javascript Object Notation (JSON) files which include metadata relating to the animal, the protocol and stimulation parameters as well as information on the host device and any other USB peripherals. Session files can additionally be exported to comma-separated value (CSV), hierarchical data format (HDF5) as well as Neurodata Without Borders (NWB) files (Rübel et al., 2019). The NWB format is a particularly useful tool in integrating behavioral and neurophysiological data in a standardized and widely accessible format. To ease the onboarding of users with Visiomode’s behavioural data, we provide an example Jupyter notebook at https://doi.org/10.5281/zenodo.6877795, which can be used as a template.

### Touchscreen behavioral arena

Next, we designed a behavioral arena that conforms to a rectangular design common across many operant chambers, measuring 200 mm (L) x 200 mm (W) x 400 mm (H).

The walls of the arena were constructed from red transparent acrylic panels (ER Plastics, UK), allowing for easy monitoring of animal behavior, while minimizing confounding external visual stimuli. Individual panels were mounted on aluminum struts (RS Components, UK) creating the exterior shape of the arena and enabling rapid assembly and disassembly during cleaning. The touchscreen (Hyperpixel 4.0, 58 mm x 97 mm, Pimoroni, RS Components, UK) was positioned on one side of the arena, accessed via a 90 mm X 115 mm hole cut in the back wall of the arena and mounted vertically across the GPIO ports of a Raspberry Pi computer (Revision 4, RS Components). To limit access to the touchscreen, a clear, transparent acyclic panel (100 mm x 125 mm x 2mm) was positioned directly in front of the screen using magnetic mounting strips attached to the arena wall. For nose poke experiments we used a panel with a 40 mm x 20 mm cut out, positioned 10 mm above the floor of the arena, while for forelimb reaching, we designed a divider with two 35 mm x 4 mm vertical slits positioned 15 mm apart (Figure 3a-b). The narrow width of the slits and position of the screen 6 mm from the arena wall prevents screen touches using the snout or tongue.

**Figure 3.**
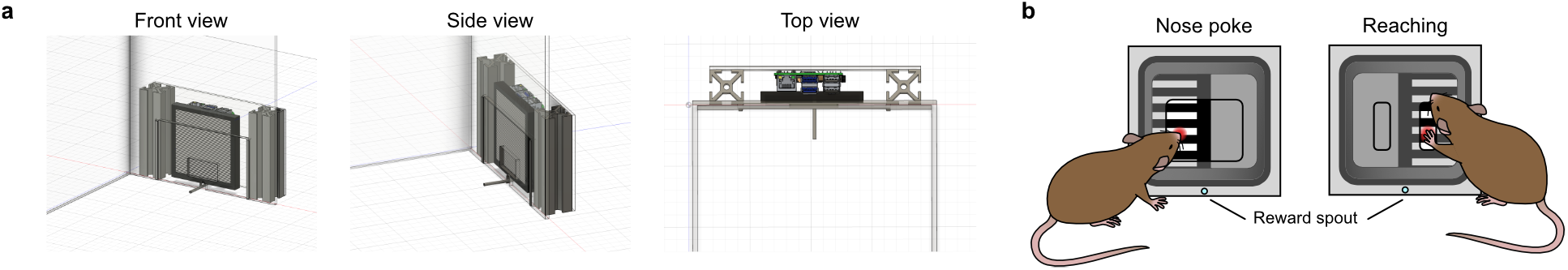
Behavioral arena with touchscreen module. (a) 3D reconstruction of the behavioral arena and touchscreen module consisting of a Hyperpixel Display 4.0, servo controlled reward spout, servo motor, solenoid, water reservoir and transparent Perspex screen divider with either nose poke or reaching slit cutouts. The behavioral arena and touchscreen module can be custom designed to fit the needs of each individual experiment. (b) Schematic diagrams showing 2-AFC nose poke (*left*) and forelimb reaching (*right*) configurations. Note the reward spout retracts after each trial using a servomotor which rotates by 10 degrees.

For mouse behavior, the size of the capacitance touchscreen is in general inversely proportional to the sensitivity, so care must be taken to select a touchscreen with a sensitivity range that is compatible with mouse touch pressures. A variety of off-the-shelf touch-screens are available for the Raspberry Pi, however, we found that the Hyperpixel display offers a good compromise between size and sensitivity. Visiomode interfaces with touchscreen devices connected to its Raspberry Pi host via the host’s operating system. This allows for Visiomode to work with the wide range of touchscreen devices with no additional configuration. Consequently, touchscreen calibration, which should not be required in most instances, would not be performed within the Visiomode interface but rather via the operating system’s own tools. To use touchscreen devices for which the Raspberry Pi operating system offers no out-of-the-box solution for calibration, users should, in the first instance, refer to the manufacturer’s specification for device drivers compatible with the Linux kernel. Alternatively, Visiomode supports the integration of custom-written hardware drivers through its devices interface, which allows for the custom integration of any external touchscreen device.

To record the behavior of the mouse in the arena (e.g., open field locomotion, rearing, grooming and task engagement) a webcam (Logitech, UK) was mounted on the wall of the arena opposite the touchscreen using a custom 3D printed support arm (https://doi.org/10.5281/zenodo.6877081).

Upon successful completion of a trial, we delivered a water reward via a 2 mm diameter clear acrylic spout mounted directly below the touchscreen. The reward spout was gravity fed via a water reservoir positioned 30 cm above the arena and dispensation was controlled by a 5 V solenoid valve (RS Components, UK) connected to an Arduino Nano microcontroller (RS Components, UK). Solenoid opening times and the height of the reservoir were adjusted to reproducibly release 10 μl of water per rewarded trial (measured using a calibrated pipette). To prevent mice from chewing or pulling the spout after reward delivery it was mounted to a 5 V servomotor which rotated the spout by 10 degrees after a 1 s delay effectively retracting the spout. While we used a solenoid valve to dispense water, the reward mechanisms could be replaced by a syringe pump allowing for finer and dynamic reward size control (Amarante et al., 2019), or by an automated food hopper mounted to the side of the touchscreen module (Acosta-Rodríguez et al., 2017). Full behavioral arena and touchscreen module build details can be found at https://doi.org/10.5281/zenodo.6877081.

### Animals, habituation, and water control

For all behavioral experiments, male adult C57BL/6J wild-type mice (8-10 weeks old, 20-30g, 3-4 animals per cage) were maintained on a reversed 12:12 hour light:dark cycle and provided ad libitum access to food and water as well as environmental enrichment. All experiments and procedures were approved by the University of Edinburgh local ethical review committee and performed under license from the UK Home Office in accordance with the Animal (Scientific Procedures) Act 1986. Mice were handled extensively for at least 5 days prior to any behavioral training and were trained once per day for 30 mins. To increase task engagement, mice were placed on a water control regime (1 ml / day) and weighed daily to ensure body weight remained above 80% of baseline (Dacre et al., 2021).

### Data Analysis & Statistics

Data were analyzed using custom scripts written in Python v3.7. Data are reported as mean ±95% bootstrapped confidence interval (95% CI) (10,000 bootstrap samples, 50 replicates per sample) unless otherwise stated. To assess the ability of mice to discriminate between the two visual stimuli in the 2AFC task, we calculated a discriminability index (d’), defined as

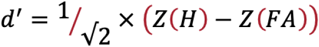

where Z(x) is the inverse-normal transformation of x, and H and FA correspond to the hit and false alarm rates, respectively (Stanislaw & Todorov, 1999). All analysis is available in the form of Jupyter notebooks at https://doi.org/10.5281/zenodo.6877795.

## Results

### Phase I: simple stimulus-response task

In the first phase of training mice had to learn a simple stimulus-response behavior. Each trial began with a fixed-length (3 s) inter-trial -interval (ITI), where the touchscreen was left blank (default is a solid black screen), before presentation of a 10 s drifting grating stimulus (100% contrast, 1 Hz sinusoid at 30 cycles / degree). During this shaping phase, mice explored the behavioral arena and responded to the presentation of a drifting grating by nose poking the touchscreen to receive a 10 μl water reward. The touchscreen rests behind a clear, transparent insert which restricts the touchable surface thus centralizing contact points. Any contact with the screen during the ITI was deemed a ‘uncued touch’ and resulted in a reset and commencement of a subsequent ITI. To allow time for reward consumption during successful trials we implemented a 1 s delay prior to the start of the next trial (Figure 4a-b). Mice rapidly learned the association between presentation of the target stimulus, response and reward displaying an increased number of successful trials and fewer miss trials (i.e., no touch response during stimulus) (Figure 4c-d). Miss trials and uncued touches were not punished. On average, mice required two behavioral sessions to reach our experimenter-defined threshold of >70 successful trials for 2 consecutive days, before they transferred to Phase II of the behavior (mean = 1.9 days [1.7, 2.1] 95% CI, N = 9 mice) (Figure 4e).

**Figure 4.**
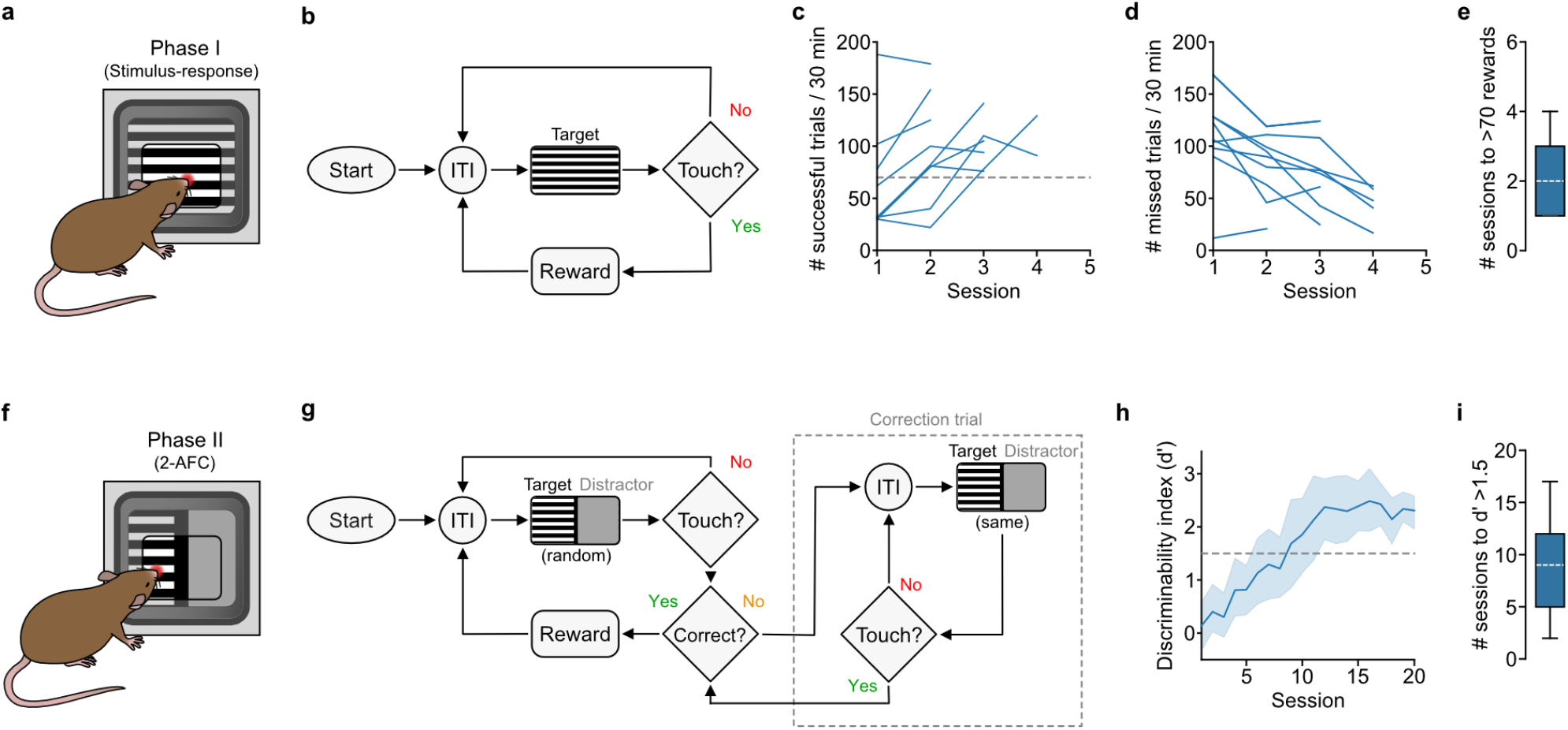
Using Visiomode to shape stimulus-response associations and 2-alternative forced choice task learning. (a) Schematic showing a mouse engaged in a simple stimulus-response behavior (Phase I). (b) Training flowchart showing Phase I of the behavioral task (simple stimulus-response association): presentation of a target stimulus (moving grating) after a fixed-length inter-trial-interval (3s, ITI) requires mice to nose poke the touchscreen to receive a water reward, failure to touch the screen results in the initiation of a subsequent ITI. (c) Number of successful trials per 30 minute training session (blue lines, data from individual mice, N = 9 mice). Gray dotted line, threshold of >70 rewards / session. Mice progress to Phase II after achieving >70 rewards / session for two consecutive sessions. (d) Number of missed trials per 30 minute training session (blue lines, data from individual mice, N = 9 mice). Note, decrease in the number of missed trials is reflected in the increase in the number of successful trials shown in (c). (e) Box-and-whisker plot showing median, interquartile range, and range of the number of sessions required to reach >70 rewards (N = 9 mice). (f) Schematic showing a mouse engaged in a 2-AFC nose-poke task (Phase II). (g) Training flowchart showing Phase II of the behavioral task (2-AFC): presentation of a pair of stimuli (target stimulus = moving grating; distractor stimulus = isoluminescent gray screen) after a pseudo-random inter-trial-interval (4-6s, ITI) requires mice to nose poke the target area of the touchscreen to receive a water reward, failure to touch the screen results in the initiation of a subsequent ITI. Nose poking the distractor area of the touchscreen results in a correction trial, where the same pair or stimuli are presented until a correct response has been achieved. (h) Discriminability index (d’) as a function of the number of training sessions. Blue line, average d’ ± 95% CI (N = 9 mice). Gray dotted line, d’ threshold of 1.5. (i) Box-and-whisker plot showing median, interquartile range, and range of the number of training sessions required to reach d’ > 1.5 (N = 9 mice).

### Phase II: 2-AFC visual discrimination nose poke task

After successful completion of behavioral shaping in Phase I, mice progressed to the 2-alternative forced choice (2-AFC) version of the task which incorporated a pseudo-randomized (4-6 s) ITI, with the target stimulus (drifting grating, 10 s) presented alongside an isoluminant grey distractor stimulus. The 10 mm ‘dead zone’ separating the stimulus and distractor was an area where touch events would not be registered (black vertical line, Figure 4f). The left versus right positioning of the target and distractor stimuli were pseudo-randomly ordered per trial. Mice learned to nose poke the target stimulus to receive a 10 μl water reward, while ignoring the distractor stimulus. If mice incorrectly chose the distractor stimulus, the subsequent trial was a correction trial, whereby the same stimulus placement was presented until the mouse correctly touched the target stimulus (Figure 4g). Mice rapidly learned to discriminate between target and distractor stimuli, with reliable discrimination (discrimination index d’ > 1.5) after an average of 9 training sessions (mean = 8.7, [7.4, 10.0] 95% CI, N = 9 mice) with peak discrimination reflecting very high performance (d’ mean = 2.4 [2.2, 2.6], N = 9 mice) (Figure 5h-i). To ensure an unbiased measure of discriminability, correction trials were excluded from d’ calculations. In addition, task engagement (i.e., hit trials / total trials) was consistently high across behavioral sessions (bootstrap mean = 92.3% [84.3, 100.0] 95% CI response rate across trials, N = 9 mice) despite mice routinely receiving more than their daily allowance of water during the task (cumulative volume of rewards > 1 ml per session) (data not shown). After ∼12 sessions, d’ became asymptotic and at 20 sessions mice were transferred to Phase III of the task.

**Figure 5.**
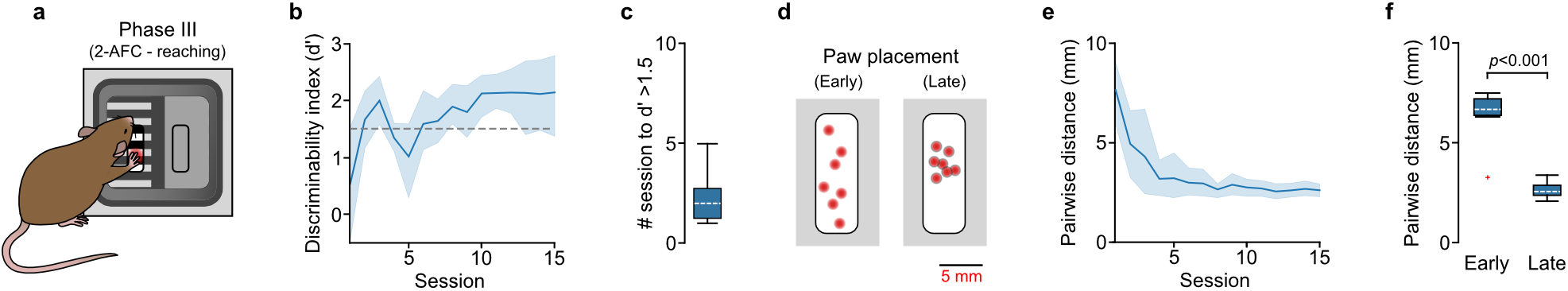
A 2-AFC visual discrimination reaching task for freely moving mice. (a) Schematic showing a mouse engaged in a 2-AFC reaching task (Phase III). (b) Discriminability index (d’) as a function of the number of training sessions. Blue line, average d’ ± 95% CI (N = 9 mice). Gray dotted line, d’ threshold of 1.5. (c) Box-and-whisker plot showing median, interquartile range, and range of the number of training sessions required to reach d’ > 1.5 (N = 9 mice). (d) Schematic showing paw placement distributions during an early (*left*) and late (*right*) training session. Red dots depict individual paw placements within the open slit. (e) Pairwise distance between individual paw placements as a function of the number of training sessions. Blue line, average pairwise distance (mm) ± 95% CI (N = 9 mice). (f) Box-and-whisker plot showing median, interquartile range, and range of the pairwise distance between individual paw placements during an early and late training session (N = 9 mice). Red cross denotes identified outlier. *p* = 2.5×10^−182^.

### Phase III: 2-AFC visual discrimination reaching task

To explore the use of forelimb reaching as an alternative readout in our 2-AFC task, we replaced the transparent ‘nose poke’ insert with an insert containing two guide slits spaced 15 mm apart. We encouraged reaching by placing the touchscreen 8 mm from the front of the slit (2 mm deep insert + 6 mm space between insert and touchscreen) which restricted access such that mice could neither nose poke nor contact the screen using their tongue (Figure 5a). Mice had to learn to reach through the slit corresponding to the target stimulus to gain a 10 μl water reward, while ignoring the slit associated with the distractor stimulus. This task setup allows investigation of sensory perception, decision-making and skilled motor control with quantifiable metrics for each component. Given the increase in complexity of the movement when switching from nose poke to reaching, mice inevitably made more mistakes resulting in a reduction in d’ immediately after the transition. This was not due to inactivity as mice displayed many cue-triggered reaches but with a high proportion of misses (mean = 168.8 reaches [160.9, 176.6] 95% CI, N = 9 mice), comparable to the number of nose pokes in Phase II (mean = 180.5 nose pokes [169.3, 191.8] 95% CI; median difference = 16.6 responses [-9.7, 34.6] 95% CI, p = 0.14; N = 9 mice).

Mice also explored using nose pokes and licking as a strategy before associating reaching with reward. The exploration phase lasted for only a short period of time with d’ recovering to > 1.5 within 2 training sessions (mean = 2.3 days [2.0, 2.7] 95% CI, N = 9 mice) (Figure 5b-c). A hallmark of rodent motor learning is the development of reproducible, stereotyped reach trajectories (Becker and Person, 2019; Galinanes et al., 2018; Kawai et al., 2015). By comparing the average pairwise distance between paw touch positions across learning, we could demonstrate the rapid decrease in pairwise distance across training, resulting in highly clustered touch positions after ∼10 training sessions (early, mean = 6.3 [5.9, 6.7] 95% CI; late, mean = 2.6 [2.5, 2.8] 95% CI; median difference = 4.1 [3.8, 4.4] 96% CI, p = 2.5×10^−128^, N = 9 mice) (Figure 5d-f). Together, our results show that mice rapidly accommodate the switch in task structure, transferring from nose poke to visually guided reaching with minimal extra training. To facilitate uptake, we have generated a detailed, step-by-step protocol for behavioral training that can be found at https://dx.doi.org/10.17504/protocols.io.bumgnu3w.

## Discussion

Here we have developed Visiomode (www.visiomode.org), a complete open-source software and hardware platform for building touchscreen-based behavioral tasks for rodents. As a key design principle, our aim was to develop a platform that was low cost and as close to turnkey as possible without sacrificing flexibility.

Visiomode’s goal is to empower users with little or no programming experience to run their own touchscreen tasks without the up-front cost of a commercial solution. After setting up a behavioral arena to the specifications we describe in this paper, the typical Visiomode user is four clicks away from their own battery of touchscreen-based behavioral tasks. First, a user would navigate to our website at www.visiomode.org, download and install the software with the instructions provided, choose from a wide selection of pre-programmed task paradigms, and click start. In contrast with currently available open-source solutions (Gurley, 2019; O’Leary et al., 2018; Pineno, 2014; Buscher et al 2020), Visiomode is openly available online and with no programming experience required due to its web interface that encapsulates all the functionality required to design tasks, as well as acquire and export behavioural data. The platform can be parallelized with no additional effort; users can follow the same steps for adding additional arenas, all of which can then be controlled from the same web browser. Thus, Visiomode is a unique, turkey solution which addresses many of the shortcomings of currently available open-source touchscreen solutions (i.e. ease of use, paradigm flexibility, code accessibility), and addresses many of the unmet needs of the user community (i.e. easy parallelization, low cost, adaptable).

Visiomode has been designed to be a community-driven project. Project development takes place on GitHub with a transparent development roadmap (https://github.com/DuguidLab/visiomode). Members of the community can contribute plugins for new protocols, stimuli and external USB devices, report bugs or suggest improvements in the software, help with documenting Visiomode or contribute to the development of the core API. The project will follow the “fork and pull” model of open-source software development for contributions, whereby any user can obtain their own copy of the source code to make changes, avoiding the need for change-related permissions. Submitted changes will be audited by the host lab to ensure all code conforms to the core project style. The “fork and pull” model empowers any user to become a contributor by eliminating the need for individual contributions to be coordinated by the project’s core team, while the review process ensures Visiomode’s stability and compatibility across behaviors. Our vision is that community-based contributions (protocols, stimuli etc) will eventually generate a comprehensive ‘go-to’ solution for any form of touchscreen-based behavior.

The Visiomode platform can accommodate a wide range of hardware configurations, including the addition of multiple USB-connected peripherals acting as input or output devices and various sized touchscreen to suit both rat and mouse behavioral arenas (Arulsamy et al., 2019; Brasted et al., 2002; Brigman et al., 2010; Bussey et al., 1998; Bussey et al., 2008; Delotterie et al., 2014; Glover et al., 2020; Gurley, 2019; Haddad et al., 2021; Heath et al., 2019; O’Leary et al., 2018; Odland et al., 2021; Piantadosi et al., 2019; Pineno, 2014; Stirman et al., 2016; Talpos et al., 2008). In addition, it permits rapid scaling to parallelize multiple behavioral arenas, each hosting its own web interface that is accessible via any personal device web browser connected to the same network. Unlike parallelized behavioral setups that are controlled by a single GUI running via a centralized host, Visiomode’s decentralized web interface design allows for multiple behavioral setups running separate tasks to be controlled in parallel. This parallelization model will facilitate high-throughput behavioral testing, enabling large-scale behavioral assays, efficient drug screening, or disease model phenotyping. This combined with the easy-to-use USB plug-and-play design permits rapid scaling up and scaling down of experiments on-the-fly depending on experimental requirements.

By developing an open-source, rapidly scalable and low-cost platform our aim is to increase uptake of touchscreen-based behavioral assays across our community, accelerating the investigation of cognition, decision-making and sensorimotor behaviors both in health and disease.

## Author Contributions

Conceptualization and design, C.E., and I.D.; methodology & investigation, C.E., V.P., T.C., and I.D.; resources, C.E., T.C.; graphic design, M.Z.; writing, reviewing & editing, all authors.

## Data Availability

All code and data are available at https://github.com/DuguidLab/visiomode/.

## Acknowledgment

We are grateful to members of the Duguid lab for experimental discussions and comments on the manuscript. This study was supported by the Simons Initiative for the Developing Brain PhD studentship (C.E.) and a Wellcome Senior Research Fellowship (UK) (110131/Z/ 15/Z) to I.D.

